# The Effect of A1/A2 Reactive Astrocyte Expression on Hydrocephalus Shunt Failure

**DOI:** 10.1101/2021.11.04.467357

**Authors:** Fatemeh Khodadadei, Rooshan Arshad, Diego M. Morales, Jacob Gluski, Neena I. Marupudi, James P. McAllister, David D. Limbrick, Carolyn A. Harris

## Abstract

Understanding the composition characteristics of the glial scar contributing to the high failure rate of neuroprosthetic devices implanted in the brain has been limited, to date, with the evaluation of cells, tissue, and biomarkers obstructing the implant. However, there remains a critical knowledge gap in gene expression profiles of the obstructing cells. This first-time study investigates the phenotypic expression specific to astrocyte scarring from those cells on hydrocephalus shunt surfaces at the time of failure, aimed at the development of therapeutic approaches to target reactive astrocytes for improved functional outcome. Recent evidence has indicated that the tissue obstructing shunts is over 80% inflammatory, with a more exaggerated astrocytic response. To understand how to mitigate the astrocyte immune response to shunts, we performed gene expression profiling of the C3 and EMP1 genes to quantify if astrocytes were classically activated and pro-inflammatory (A1) or alternatively activated and anti-inflammatory (A2), respectively. Shunt catheters were removed from patients at the time of failure and categorized by obstructed vs non-obstructed shunts. RNAscope fluorescent in situ hybridization and quantitative PCR analysis of the C3 and EMP1 expressed genes revealed that a heterogeneous mixed population of both the A1 and A2 reactive phenotype exist on the shunt surface. However, the number of A2 reactive astrocytes are significantly higher on obstructed shunts compared to A1 reactive astrocytes. ELISA data also confirmed higher levels of IL-6 for obstructed shunts involved in A2 reactive astrocyte proliferation and glial scar formation on the shunt surface. Since TNF-α and IL-1β propel resting astrocytes into an A2 reactive state, by simply blocking the secretion or action of these cytokines, astrocyte activation and attachment on obstructing shunts could be inhibited.

## 1. Introduction

Implantation of foreign materials within the brain initiates a series of reactions, collectively called the foreign body reaction (FBR), which aims to eliminate or isolate the implanted foreign material from the host immune system. Upon implantation of large medical devices such as neuroprosthetics, where elimination is not possible, the FBR continues until the device is barricaded from healthy brain tissue. The initial phase of the FBR is blood-device interactions, which occurs immediately upon implantation caused by vasculature or blood-brain barrier (BBB) disruption. This results in the nonspecific adsorption of blood proteins to the device surface through a thermodynamically driven process to reduce surface energy. Other than BBB disruption and influx of serum proteins, the immune system is also activated by signals of host cell injury and extracellular matrix (ECM) breakdown proteins such as fibrinogen and fibronectin adhesion to the device surface. Microglia, the resident immune cells of the central nervous system (CNS), and blood-derived macrophages recognize the protein signals through receptor-mediated pathways such as toll like receptors (TLRs). Ligand binding to TLRs leads to activation of microglia/macrophages and the secretion of pro-inflammatory cytokines such as TNF-α, IL-1α, and IL-1ß [1], [2]. These very potent signaling molecules are rapidly upregulated in the injured CNS, and are observed right at the device-tissue interface corresponding to the location of activated microglia/macrophages and exaggerated astrocytes [2]–[6]. The effect of TNF-α and IL-1β is strongest on astrocyte activation and proliferation, the key member of the CNS immune response. Reactive astrocytes form a physical barrier, known as glial scar, where newly formed and hypertrophic astrocytes overlap and play a beneficial role to prevent injury from spreading to surrounding healthy tissue. However, in relation to its effect on implants, the glial scar is considered undesirable because, regardless of the device type, elicits failure [7], [8]. Collectively, the dominant role of cytokines in orchestrating the dynamic crosstalk among cells and mediating device failure is evident.

A deeper understanding of astrocyte phenotype leads to a more accurate interpretation of failure in chronically indwelling neuroprosthetics. In a recent world-renowned study, Barres and colleagues revealed two significantly different reactive astrocyte phenotype, A1 and A2 [9], [10]. The A1 reactive astrocytes produce large volumes of pro-inflammatory substances and neurotoxin that can induce neuronal death. The A2 reactive astrocytes upregulate anti-inflammatory substances and many neurotrophic factors, which promote survival and growth of neurons. The A1 neuroinflammatory astrocytes are induced by NF-κB signaling, whereas the A2 scar-forming, proliferative astrocytes are induced by STAT3-mediated signaling [9], [11]. Since glial scar borders are formed by newly proliferated, elongated astrocytes via STAT3-dependent methods, studies strongly suggest that the A2 reactive astrocyte phenotype is present during glial scar formation [10], [12], [13]. Furthermore, in vivo quiescent astrocytes that contact serum upon injury and BBB disruption, express many of the A2 reactive astrocyte genes [11], [14], [15].

In the brain, TNF-α, IL-1α and C1q combined propel resting astrocytes into an A1 reactive state [9]. Co-stimulation with TNF-α and IL-1β induces the A2 reactive state with neurosupportive characteristics [16]. In fact, TNF-α and IL-1β modulate the glial scar process by stimulating astrocyte IL-6 secretion [17]. IL-6 primarily activates astrocyte proliferation by a positive feed-forward loop, further activating local astrocytes to maintain the glial scar through self-sustaining mechanisms. IL-6 signaling pathways, are enhanced in A2 reactive astrocytes, and STAT3 is activated by IL-6 [11], [18]. IL-6 is one of the initial triggers of reactive astrocytes in the acute phase of disease, involved in improving neuronal survival and neurite growth [7], [8]. Although, these properties are evidence of the beneficial roles of IL-6 in repair and modulation of inflammation in the CNS, the overproduction of IL-6 is associated with glial scar formation. Hence, a careful inflammatory balance of IL-6 is essential for proper repair. Inhibition of both IL-6 and IL-6r by antibody neutralization reduces glial scar formation on the implanted device and damage to the brain as a result of bystander effects of increased CSF cytokine levels [19].

Hydrocephalus is a devastating and costly disease. The most common treatment paradigm is surgical shunting of cerebrospinal fluid (CSF). However, shunts are plagued by unacceptably high failure rates (40% in the first year, 90% in the first ten years) [20]–[23], and impose a significant burden on patients, their families, and society. Understanding the root causes of shunt failure to design improved devices, will indeed reduce this burden. Shunts primarily fail due to obstruction of the shunt system with adherent inflammatory cells [24]–[29]. Astrocytes and macrophages are the dominant cell types bound directly to the catheter. Our recent work indicates that astrocytes make up more than 21% of cells bound to obstructed shunts. We have also observed that, of the obstructed masses blocking ventricular catheter holes, astrocytes makeup a vast majority of cells. We have also found astrocyte markers in obstructive masses to be co-localized with proliferative markers, indicating that astrocytes are active on the shunt surface; they produce inflammatory cytokine IL-6 and proliferate [30], and their number and reactivity peak on failed shunts. In hydrocephalus patients, IL-6 cytokines significantly increase during shunt failure, especially after repeat failures. Astrocytes create a “glue” for more glia or other cells and tissues to secondarily bind and block the shunt. Even contact with the ventricular wall results in astrocyte migration to the surface and interaction with the shunt [31].

In this study, our goal is to reduce shunt obstruction by re-developing a strategy to reduce astrocyte activation and thus attachment and density on the shunt surface. Thus, we must control the degree of inflammatory cell activation. This is not the direct target of a specific treatment paradigm to date. We must first determine astrocyte phenotype expression in tissue that is obstructing shunts using failed patient’s shunts, then employ a promising pharmacological agent that will inhibit the cell activation state to reduce attachment. The A1 reactive astrocytes are not as quickly proliferative, however, considerable proliferation of A2 reactive astrocytes is seen when the reactive response is to produce a protective scar around the injury [10]. Our goal is to observe whether the cells blocking shunts are expressing an A1 or A2 reactive astrocyte phenotype to understand how to mitigate the cell immune response to shunts. That is to reduce mechanisms leading to shunt failure through inhibition of cell activation. To address this new research avenue, we use RNAscope fluorescent in situ hybridization and quantitative PCR analysis to determine the A1 or A2 reactive astrocyte phenotype expression on failed shunt. ELISA analysis confirms the pro- and anti-inflammatory cytokine concentration profiles in the CSF associated with astrocyte activation. A powerful marker for A1 is the classical complement cascade component C3, specifically upregulated in A1 reactive astrocytes (and not in resting or A2 reactive astrocytes). C3 is one of the most characteristic and highly upregulated genes in A1 and EMP1 is an A2-specific gene. Then we employ the release of an FDA-approved pharmacological agent on the shunt surface that will inhibit the cell activation state to reduce the presence of astrocytes on shunts. This will keep any attaching astrocytes in a resting state, reduce proliferation, inhibit downstream proliferation, and ultimately deter shunt obstruction. Since the master cytokine IL-1 (α and β) is the initial molecular mediator that triggers glial scar formation around other devices in the brain, we will investigate whether astrocytes obstructing shunts could be prevented by simply blocking secretion or action of these cytokines to keep astrocytes out of the A1 or A2 reactive state. FDA-approved drugs targeting TNF-α, IL-1α and IL-1β already exist and are in use for other medical conditions.

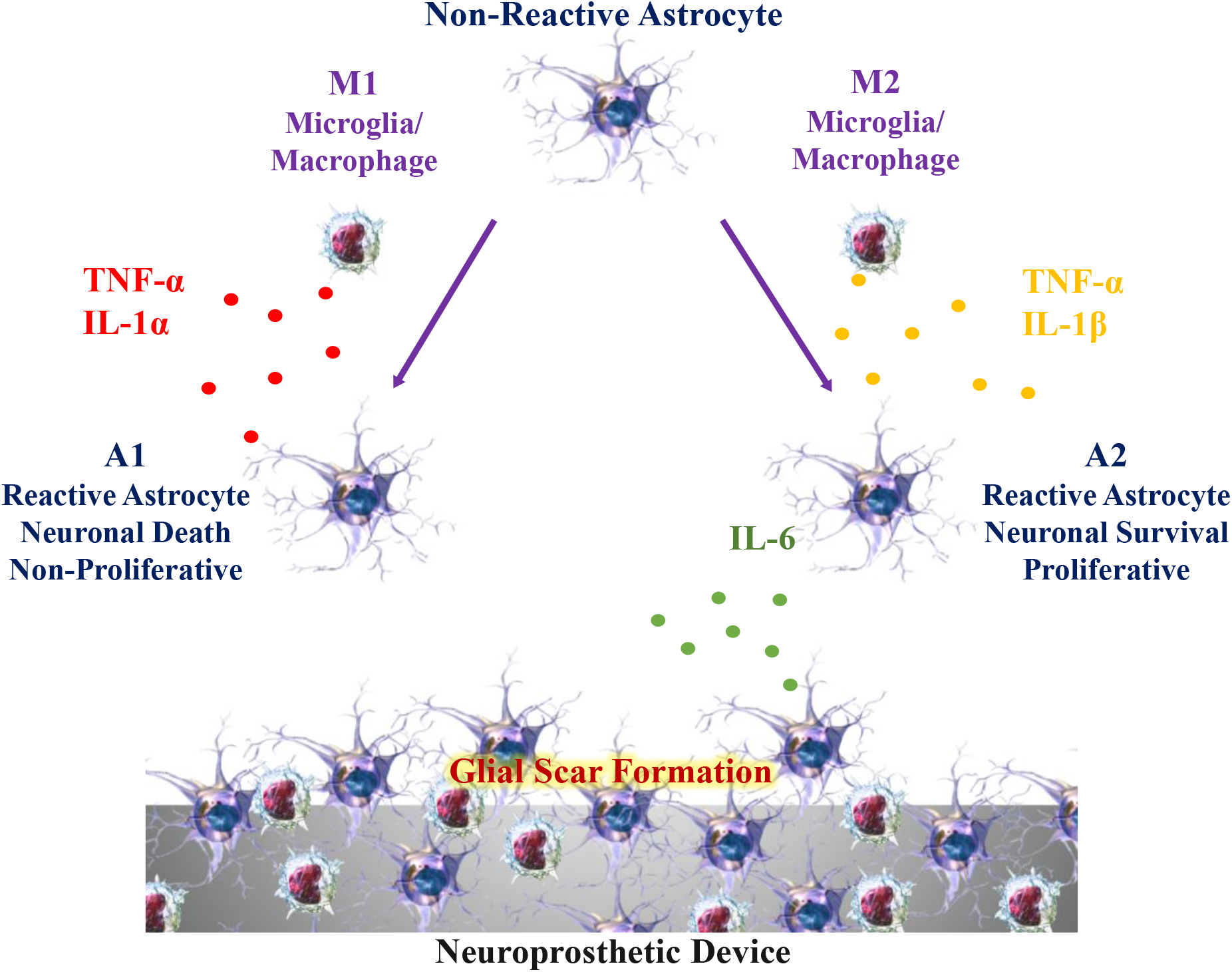
Schematic. Microglia/macrophage and astrocyte reactions following neuroprothetic device implantation. Injury transforms microglia into an M1- and M2-like phenotype and astrocytes into an A1- and A2-type, correspondingly. Astrocytes and microglia work together to initiate either a neuroinflammatory or neuroprotective response after injury through the release of cytokines or neurotrophic factors that can lead to neuronal death or survival. The cytokine pathway is the best and most important measurable outcome for inflammatory cascades. Inflammatory cells at the brain-device interface communicate via cytokines to activate and recruit other inflammatory cells to the interface. Cytokine stimulation is a gateway for other gene products to be over- or under-expressed in the cascade, resulting in device failure. Therefore, the cytokine pathway is a starting point for mechanistic, thorough investigation of inflammation and device failure.

## 2. Materials and methods

### 2.1. Ethics approval

The permission to collect shunt hardware, CSF, and patient data was approved by the Wayne State Institutional Review Board (IRB) as the coordinating center and as a participating site. Written informed consent was obtained from all patients or their legally authorized representative. Collection was performed in a manner consistent with the standard of treatment; decision to remove the shunt was always based on clinical symptoms for surgical intervention chosen by the neurosurgeon. Samples were collected from individuals with any hydrocephalus etiology and clinical history. After removal by a surgeon, the shunt samples were immediately submerged in a solution of paraformaldehyde (PFA) to fix cells for RNAscope fluorescent in situ hybridization experiments or RNAlater, an aqueous, non-toxic tissue storage reagent that rapidly permeates tissue to stabilize and protect the quality/quantity of cellular RNA in situ in unfrozen specimens for qPCR experiments. RNAlater eliminates the need to immediately process tissue specimens or to freeze samples in liquid nitrogen for later processing. Obstructed and non-obstructed shunts were characterized based on the degree of actual tissue blockage on the shunt surface shown in Figure 1B. A whole-mount procedure was used for all non-obstructed shunts, where the whole shunt was immersed in solution. For all obstructed shunts the tissue on the surface of the shunt was removed from the shunt. Tissue was then embedded in OCT compound for RNAscope fluorescent in situ hybridization experiments or immersed in solution for qPCR experiments. As previously described [32], CSF was collected at the time of shunt surgery and transported on ice to the Washington University Neonatal CSF Repository. Samples were then centrifuged (2500 rpm for 6 min) at room temperature, and the supernatant was aliquoted and stored in 1.5 ml polypropylene microcentrifuge tubes at −80 °C until experimental analysis.

**Figure 1.**
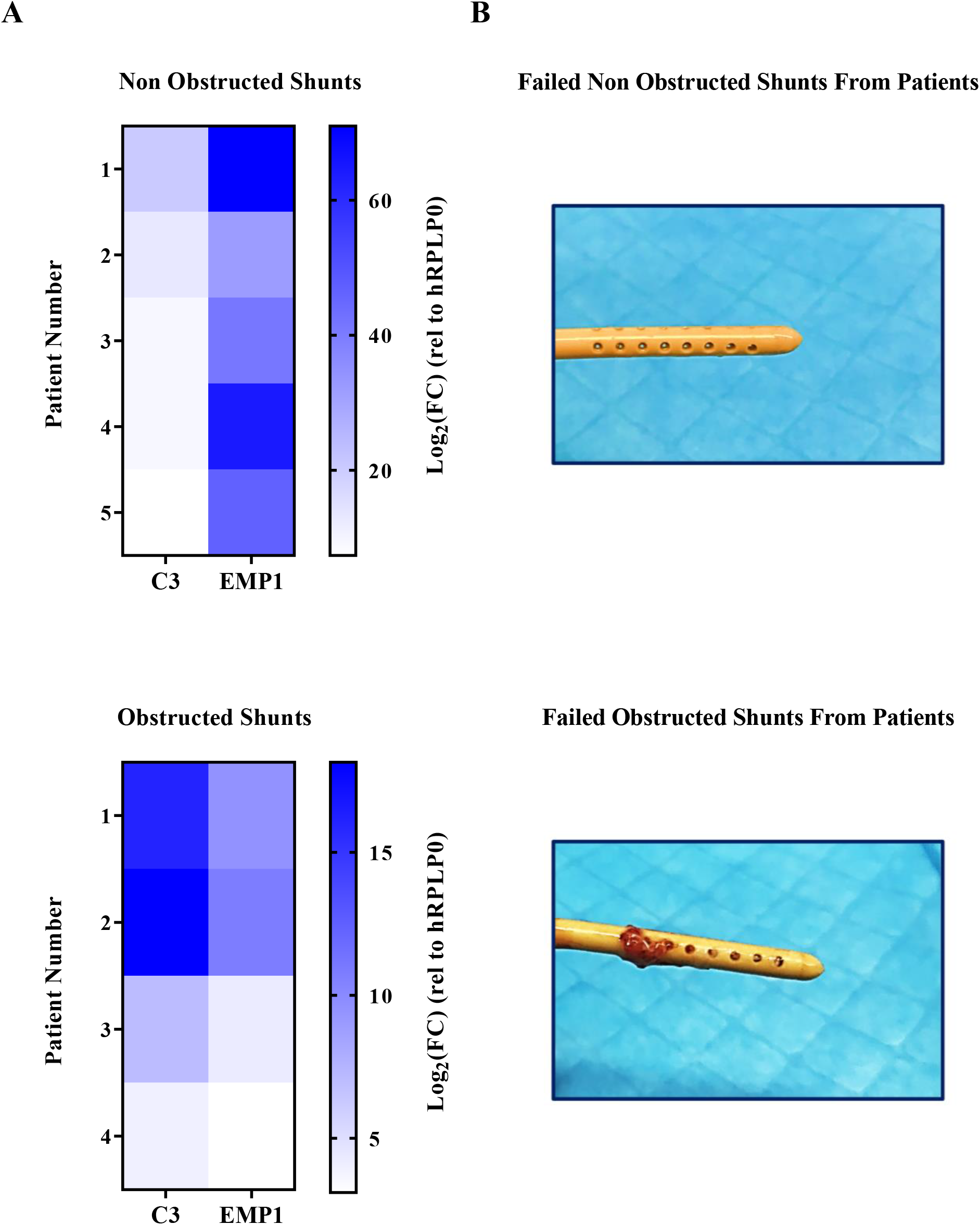
Expression of C3, EMP1 astrocyte activation genes assessed by qPCR on obstructed and non-obstructed shunts. (A) Heatmaps comparing the expression of A1-specific reactive gene C3 and the A2-specific reactive gene EMP1 for obstructed and non-obstructed shunts collected from patients (two-way ANOVA test). (B) Representative images for obstructed and non-obstructed shunts.

### 2.2. Quantitative PCR

Total RNA was extracted using the GenElute Mammalian Total RNA Miniprep Kit (sigma), cDNA synthesis was performed using the iScript cDNA Synthesis Kit (Bio-Rad), and qPCR was completed using the PowerUp SYBR Green Master Mix (Applied Biosystems) according to manufacturer protocols. Relative mRNA expression was normalized to hRPLP0 (reference gene) [33]. Primers for human are as follows: hC3 (A1 reactive astrocyte marker), hEMP1 (A2 reactive astrocyte marker) [34].

### 2.3. RNAscope fluorescent in situ hybridization

RNAscope fluorescent in situ hybridization was performed on fixed frozen tissue. Tissue was embedded in OCT compound (Tissue-Tek) and 14 μm tissue sections were prepared and immediately frozen at −80 °C. Multiplex RNAscope was performed according to manufacturer’s (ACD: Advanced Cell Diagnostics) protocol against the target mRNA probes of hC3 (label for A1 reactive astrocytes), hEMP1 (label for A2 reactive astrocytes), and hSLC1A3 (label for astrocytes). RNAscope fluorescent in situ hybridization is nonlinearly amplified and thus intensity cannot be used to measure expression. Instead, images were thresholded in ImageJ. The percent of area covered by this thresholded signal was then quantified and recorded as reactivity [33]. Images were acquired with a resonance-scanning confocal microscopy (RS-G4 upright microscope, Caliber ID, Andover, MA, USA).

### 2.4. Multiplex ELISA

Multiplex assays were run by the Bursky Center for Human Immunology & Immunotherapy Programs (CHiiPs) at Washington University School of Medicine. Frozen supernatant CSF was slowly thawed and then analyzed in duplicate with multiplex immunoassay kits according to the manufacturer’s instructions for the following inflammatory cytokines: IL-1α, IL-1β, IL-6, TNF-α, IL-8, IL-4, IL-10 (Thermofisher Scientific), C3, and C1q (Millipore Sigma). Briefly, magnetic beads were added across all the wells on the plate, CSF samples and standards were then added in duplicate. Following washing steps, the detection antibody was added followed by streptavidin incubation. Beads were then resuspended with reading buffer and data were acquired on a Luminex detection system. The concentration of each analyte was calculated by plotting the expected concentration of the standards against the multiplex fluorescent immunoassay generated by each standard. A 4-parameter logistic regression was used for the best fit curve. Protein concentration is reported as pg/mL for each analyte.

### 2.5. Purification of astrocytes by immunopanning

Astrocytes were purified by immunopanning from post-natal day (P) 5 mouse brains and cultured as previously described [35]. Cerebral cortices were dissected and enzymatically digested using papain at 37 °C and 10% CO_2_. Tissue was then mechanically triturated with a serological pipette at RT to generate a single-cell suspension. The suspension was filtered and negatively panned for microglia/macrophage cells (CD45), oligodendrocyte progenitor cells (O4 hybridoma), and endothelial cells (L1) followed by positive panning for astrocyte cells (ITGB5). Astrocytes were cultured in defined, serum-free medium containing 50% neurobasal, 50% DMEM, 100 U/mL penicillin, 100 μg/mL streptomycin, 1mM sodium pyruvate, 292 μg/mL L-glutamine, 1× SATO, 5 μg/mL of N-acetyl cysteine, and 5 ng/mL HBEGF.

All animal protocols were approved by the Institutional Animal Care and Use Committee at Wayne State University (IACUC).

### 2.6. Targeted drug delivery

A1 reactive astrocytes were generated by culturing the purified astrocytes on PDMS coated tissue culture plates and then treating for 24 h with IL-1α (3 ng/ml, Sigma, I3901), TNF-α (30 ng/ml, Cell Signaling Technology, 8902SF), and C1q (400 ng/ml, MyBioSource, MBS143105). A2 reactive astrocytes were generated by culturing the purified astrocytes on PDMS coated tissue culture plates and then treating for 24 h with IL-1β (30 ng/ml, Cell Signaling Technology, 8900SF) and TNF-α (30 ng/ml, Cell Signaling Technology, 8902SF). A1 reactive astrocytes were targeted for 48 h using neutralizing antibodies to IL-1α (30 ng/ml, Abcam, ab9614), TNF-α (30 ng/ml, Cell Signaling Technology, 7321), and TGF-β (30 ng/ml, R&D Systems, 243-B3-002/CF). A2 reactive astrocytes were targeted for 48 h using neutralizing antibodies to IL-1β (30 ng/ml, Abcam, ab9722), TNF-α (30 ng/ml, Cell Signaling Technology, 7321), and IL-6 (30 ng/ml, Abcam, ab6672) [36], [37].

Polydimethylsiloxane (PDMS) coated tissue culture plates were prepared by mixing Sylgard-184 elastomer and curing agents at a ratio of 10:1 (w/v), then pouring into the plates and curing for 48 h.

### 2.7. Data presentation and statistical analysis

All data presented was performed using GraphPad Prism version 8. Two-tailed unpaired Student’s t-test and two-way ANOVA test was performed. The mean with standard error mean (for the experiments done independently with technical replicates) and standard deviation was displayed. A predetermined significance level of *P* < 0.05 was used in all statistical tests.

## 3. Results and discussion

### 3.1. Understanding the inflammatory response following shunt implantation

Quantitative PCR (qPCR) was performed on collected tissue from failed shunts received from patients. Since the tissue obstructing shunts comprises of a more exaggerated astrocytic response, we investigated the up-regulation of the A1-specific reactive gene C3 and the A2-specific reactive gene EMP1. We observed that a heterogeneous up-regulation of both the A1- and the A2-specific reactive gene exist on both obstructed and non-obstructed shunts with no significant difference. In addition, we observed an increase in the EMP1 expression on non-obstructed shunts, while an increase in the C3 expression was noticed on obstructed shunts (Figure 1).

Indeed, A1 reactive astrocytes are a major source of the classical complement cascade component C3, however, other inflammatory cells in the tissue on obstructed shunts also induce the expression of C3. The increased expression of C3 on obstructed shunts is in accordance with other studies linking persistent neuroinflammation to neurodegeneration and adverse effects on the neural circuits and decrease excitatory neuronal function [38]. This is to recruit additional immunocytes to the site and exacerbate the secondary insult response.

Astrocytes are the dominant cell type bound directly to non-obstructed shunts and play a neuroprotective role, particularly in the acute phase of injury following an immediate disruption of the blood-brain barrier (BBB). Therefore, the increased expression of EMP1 on non-obstructed shunts is in accordance with other studies [29].

### 3. 2. Understanding the astrocyte phenotype expression on implanted shunts

RNAscope fluorescent in situ hybridization was performed on collected tissue from failed shunts received from patients. In accordance with our qPCR data, we found that the majority of SLC1A3+ astrocytes express A1 (C3) and/or A2 (Emp1) markers, suggesting that astrocytes can express a combination of A1 and A2 genes on shunt surfaces. In particular, we observed that a greater number of SLC1A3+ astrocytes expressed the A2-specific gene Emp1 on both obstructed and non-obstructed shunts. Interestingly, the number of A2 reactive astrocytes are significantly larger on obstructed shunts compared to A1 reactive astrocytes (Figure 2). In our recent work, we have also observed astrocyte markers in obstructive masses to be co-localized with proliferative markers, indicating that astrocytes are active on the shunt surface: they produce inflammatory cytokine IL-6 and proliferate. Since A2 reactive astrocytes are proliferative [10], they are responsible for the glial scar formation observed on obstructed shunts. This is in accordance with other studies, indicating that glial scar borders are formed by newly proliferated, elongated astrocytes that interact to corral inflammatory and fibrotic cells via STAT3-dependent mechanisms [13], and that astrocytes in scar formation seem to be devoid of C3 upregulation [39].

**Figure 2.**
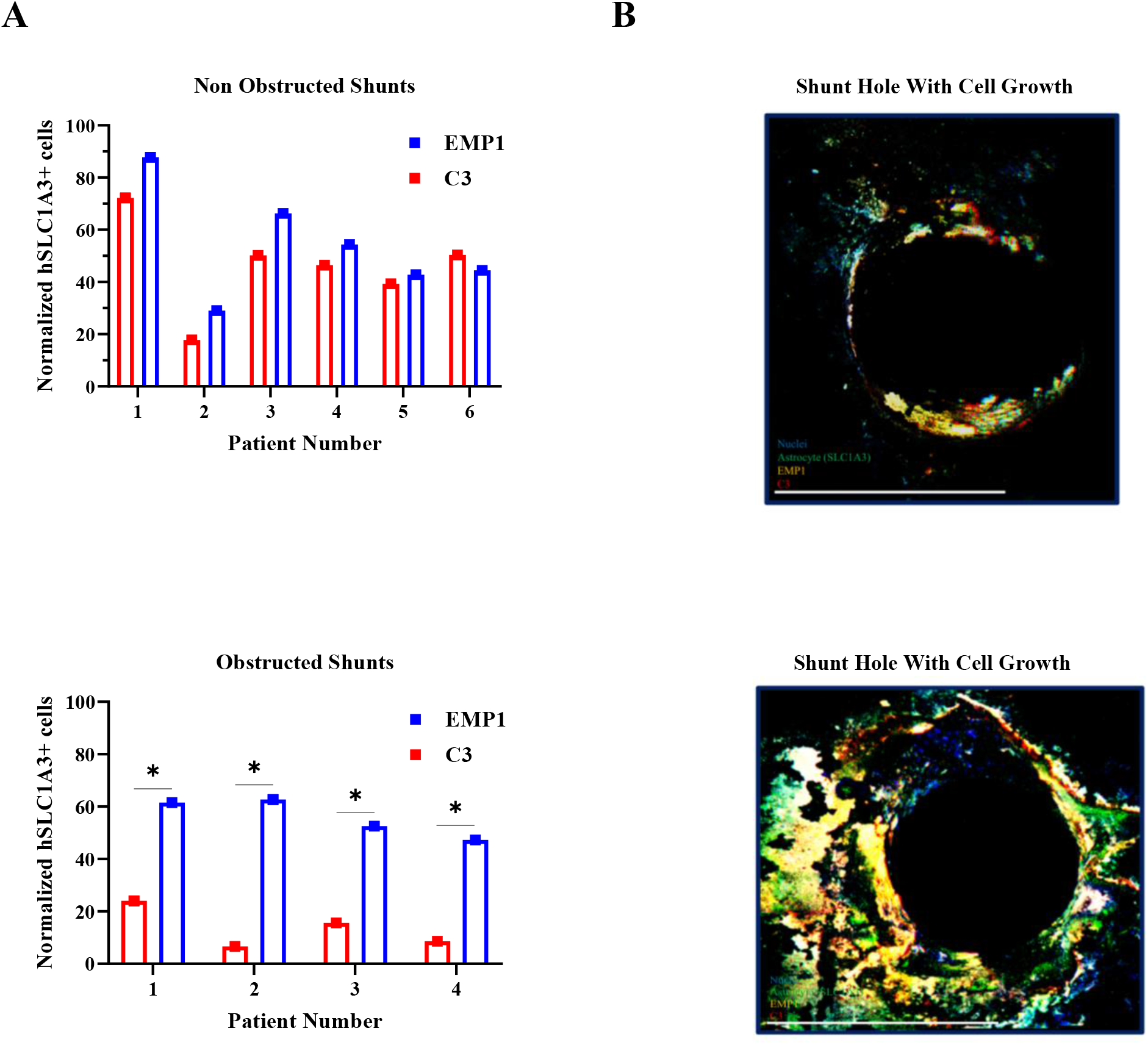
Comparison of astrocyte response by RNAscope fluorescent in situ hybridization on obstructed and non-obstructed shunts. (A) The astrocyte phenotype specificity of C3 and EMP1 RNAscope fluorescent in situ hybridization signal was assessed by probing for SLC1A3+ astrocytes on both obstructed and non-obstructed shunts collected from patients. For normalization, the C3 and EMP1 signals were dividing by SLC1A3 signals (*p < 0.05; two-way ANOVA test). (B) Representative RNAscope fluorescent in situ hybridization images for obstructed and non-obstructed astrocyte gene C3 (red) and EMP1(yellow) showing colocalization with the astrocyte marker SLC1A3 (green) (scale bar = 500 μm).

### 3.3. Cerebrospinal fluid biomarkers of neuroinflammation in obstructed and non-obstructed shunts

Using multiplex ELISA, this study investigated shunt failure through the CSF protein concentration profiles of select pro-inflammatory and anti-inflammatory cytokines for obstructed and non-obstructed shunts. C1q, IL-1α, and TNF-α induce A1 reactive astrocytes, IL-1β, TNF-α and IL-6 induce A2 reactive astrocytes. C3 is an A1 astrocyte marker. IL-8 and IL-10 inflammatory cytokines are of interest as they consistently stand out by being elevated in the CSF of hydrocephalus patients. Remarkably, in accordance with our qPCR data, we found higher neuroinflammation for obstructed shunts, however, with no significant difference compared to non-obstructed shunts confirming the heterogeneous mixed population of both the A1 and the A2 reactive astrocytes. Interestingly, higher levels of IL-6 are observed for obstructed shunts compared to non-obstructed shunts (Figure 3). As indicated in the RNAscope fluorescent in situ hybridization results, IL-6 primarily activates A2 reactive astrocyte proliferation through a positive feed-forward loop, activating local astrocytes to maintain the glial scar formation on the shunt surface.

**Figure 3.**
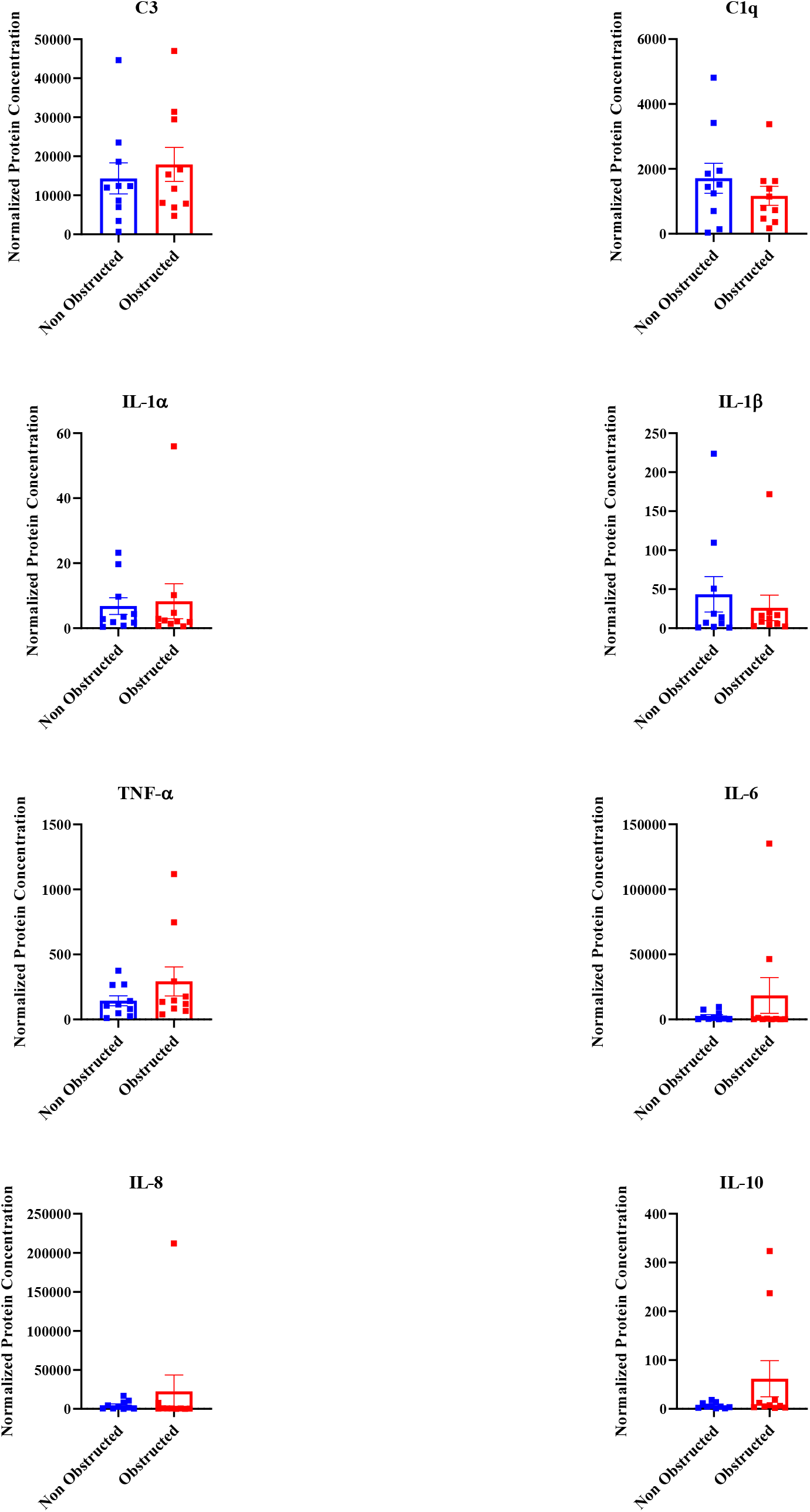
Cerebrospinal fluid cytokine concentrations between obstructed and non-obstructed shunts. Analytes include C3, C1q, and IL-1α (A1 astrocyte markers), IL-1β and IL-6 (A2 astrocyte markers), TNF-α, IL-8 and IL-10. For normalization, the concentration of each cytokine is divided by the total protein concentration for each group (two-tailed unpaired Student’s t-test, n = 10 per group, mean ± SEM).

In our recent paper, higher levels of IL-6 are observed for non-obstructed shunts compared to obstructed shunts [32]. However, obstructed and non-obstructed shunts were characterized based on the symptoms of obstruction defined by the patient’s charts instead of the degree of actual tissue blockage on the shunt surface as described in this study.

### 3.4. Inhibiting astrocyte cell activation and attachment on the shunt surface with neutralizing antibody treatment and anti-inflammatory cytokines

Now we understand that a heterogeneous mixed population of both the A1 and A2 reactive phenotype exist on the shunt surface. In addition, TNFα, IL-1α combined propel resting astrocytes into an A1 reactive phenotype [9] and co-stimulation with TNF-α and IL-1β induces an A2 reactive astrocyte phenotype [16]. Also, IL-6 induces astrogliosis and astrocyte proliferation [30]. Therefore, we investigated whether the activity of astrocytes could be significantly reduced by simply employing already FDA-approved antibody therapies that inhibit human TNF-α, IL-1α, IL-1β, and IL-6. Hence, neutralizing antibodies to TNF-α, IL-1α, IL-1β and IL-6 were employed to decrease the activity of A1 and A2 astrocytes for a significant decrease in attachment on PDMS coated surfaces mimicking the shunt surface (Figure 4). These data are in accordance with other studies indicating that the knockout of reactive astrocyte activating factors slows disease progression [33], dampening the formation of reactive astrocytes prevents neuronal death [37], and astrogliosis inhibition attenuates hydrocephalus [40].

**Figure 4.**
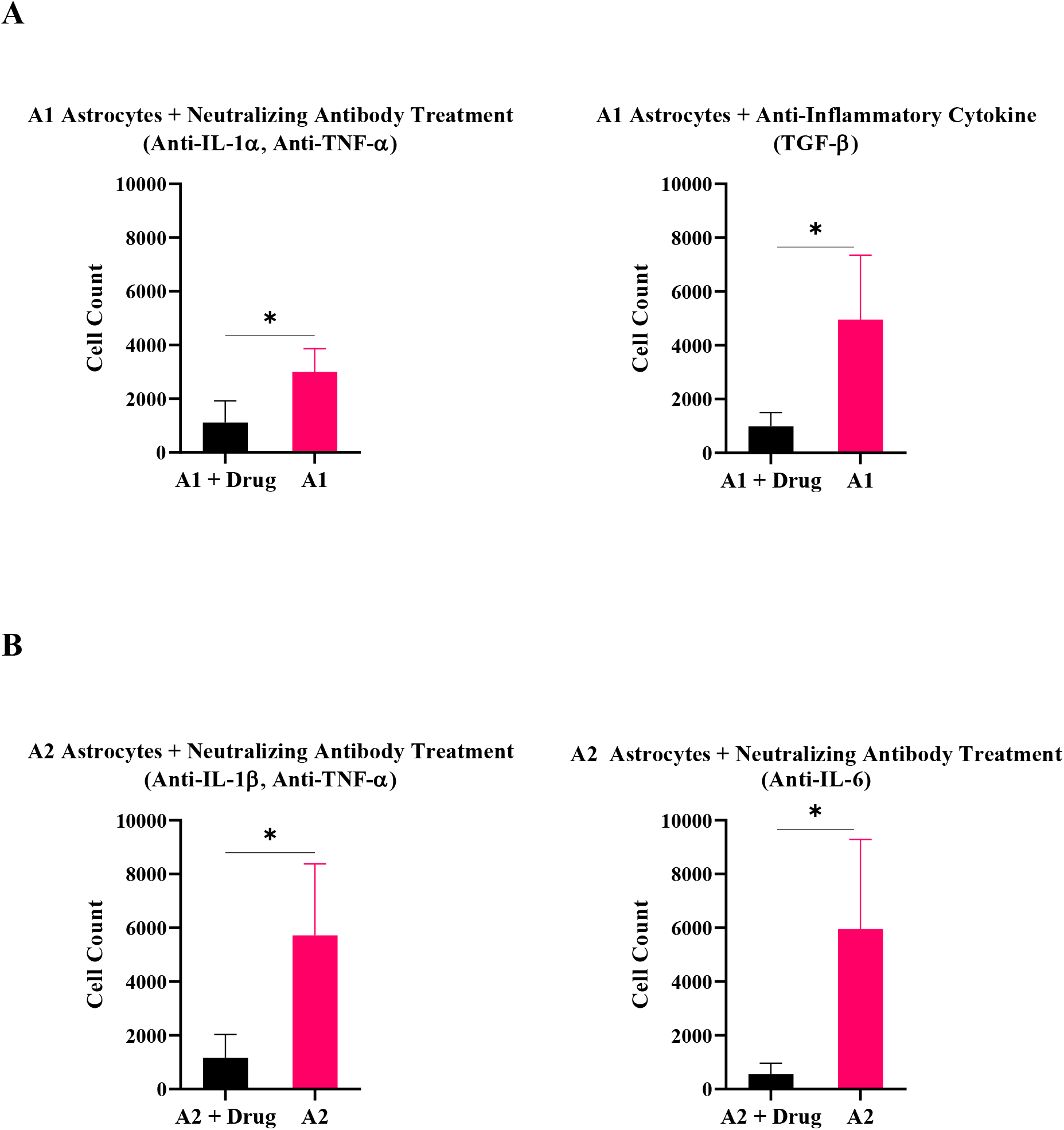
Antibody therapies that will inhibit the cell activation state to reduce attachment on the shunt surface. (A) A1 reactive astrocytes treated with neutralizing antibodies to TNF-α, IL-1α, and anti-inflammatory cytokine TGF-β. (B) A2 reactive astrocytes treated with neutralizing antibodies to TNF-α, IL-1β, and IL-6 (*p < 0.05; two-tailed unpaired Student’s t-test, n = 3 per group, mean ± SD).

The anti-inflammatory cytokine TGF-ß was able to reset A1 astrocytes to a non-reactive state, significantly reducing cell attachment on the PDMS coated surface. This is in accordance with other studies indicating that TGF-ß suppresses A1 astrocyte activation [36], reverses the formation of A1 astrocytes by fibroblast growth factor (FGF) signaling [41], and greatly reduces the expression of A1-specific markers [42]. Furthermore, TGF-β did not induce A2 reactive astrocyte attachment on the PDMS coated surface.

Taken together, these data suggest that drug therapies could eventually be added to the shunt as device coating and released in vivo for enhanced next-generation medical devices.

## 4. Discussion

This first-time study, which pulls strengths from the recent world-renowned Barres et al. study on astrocyte activation, represents a robust investigation of the changes in gene expression levels specific to astrocyte immune response following CSF shunt implantation. By shedding light on the mystery of astrocyte phenotype expression on shunt surfaces, root causes for shunt failure can be achieved to improve hydrocephalus treatment.

Cell adhesion is not necessary to drive the neuroinflammatory response and biomaterials that go beyond reducing cell adhesion alone but also incorporate improved attenuation of inflammation in the tissue surrounding the implanted device are required [6]. In extensive studies, chronically implanted neural implants with coatings were compared to that of identical uncoated devices. In vitro, the coated implant significantly reduced astrocyte and microglial adhesion by ~95% [43]. A similar reduction of cell adhesion was observed following device removal after two, four or 24 weeks of implantation in rat cortical tissue [44]. Interestingly, no significant difference was observed in the neuro-inflammatory response or the level of neuronal loss surrounding the coated implant compared to uncoated devices. A persistent inflammation was observed surrounding both uncoated and coated implants. Furthermore, neuronal density around the implanted devices was also lower for both implant groups compared to the uninjured controls. As no cells were found adhered to coated implants upon removal, both coatings were still functioning at the endpoints studied.

Our recent paper also indicates that under higher shear stress, despite less cell attachment to the surface, a significant increase in IL-6 secretion is detected [30]. Our data support others, which identify a necessity for control of the degree of inflammatory cell activation for considerable enhancement of device performance within the brain, in contrast to the common implant failure of reduced cell adhesion on the device surface in vivo [4], [45]–[47]. Our results in combination with previous studies present a proof of concept that to have significant impact, strategies should implement a focus on attenuating the initial inflammatory cell activation instead of only aiming at reducing cell adhesion on the device surface. Such strategies include decreasing shear activation as a primary cause of device failure [2], [4], [7], [8], [48]–[52], and directly antagonizing the accumulation of pro-inflammatory cytokines via targeted therapeutic for TNF-α, IL-1α, Il-1β, and IL-6.

The challenges for targeted drug delivery to A1 and A2 reactive astrocytes will be determining the parameters for delivery, specifically, the onset point, the dosage, the duration, and the delivery vehicle for anti-inflammatory device coating in vivo. Cytokine responses were strongly upregulated within a day post implantation indicating cytokine targeting strategies need to be present at the site of implantation immediately following implantation. Also, given the complexity of molecules involved in the inflammatory response, many other potential molecular targets could be identified for therapeutic purposes. However, when employing such strategies, as increasing evidence has shown that many inflammatory molecular mediators in the CNS act like double-edged swords. Their effects do not occur in an all or-none fashion; rather, they are concentration dependent. Therefore, it should be kept in mind that the goal is to contain any excessive activation rather than to remove all activity.

### Conclusion

First-time revealing of astrocyte phenotype expression leads to (1) a precise understanding of cellular response mechanism to device implantation and a precise interpretation of failure in chronically indwelling neuroprosthetics, (2) a therapeutic window for more targeted therapies to inhibit astrocyte activation and attachment on the implant. Therefore, for significant reduction in device failure, alongside manipulating shunt properties, drug therapy is used for inhibition of cytokines and therefore inhibition of cell aggregation for achieving stable and long-term functional outcomes. A combinatorial strategy that is rationally designed and tailored, aided by a better understanding of underlying biological processes, will lead to cumulative improvements in CSF shunt technology that will one day be translated into real clinical applications.

## Competing interests

The authors declare no competing interests.

## Notes

### Competing Interest Statement

The authors have declared no competing interest.

